# Niche partitioning and coexistence of two spiders of the genus *Peucetia* (Araneae, Oxyopidae) inhabiting *Trichogoniopsis adenantha* plants (Asterales, Asteraceae): a population approach

**DOI:** 10.1101/568410

**Authors:** German Antonio Villanueva-Bonilla, Suyen Safuan-Naide, Mathias Mistretta Pires, João Vasconcellos-Neto

## Abstract

Niche theory suggests that the coexistence of ecologically similar species at the same site requires some form of resource partitioning that reduces or avoids interspecific competition. Here, we investigated the temporal and spatial niche differentiation of two sympatric congeneric spiders, *Peucetia rubrolineata* and *P. flava*, inhabiting *Trichogoniopsis adenantha* (Asteraceae) plants along an altitudinal gradient in various shaded and open areas in an Atlantic forest in Serra do Japi, SP, Brazil. In theory, the coexistence of two *Peucetia* species could be explained by: (1) temporal segregation; (2) differential use of the branches of the plant; (3) differential use of specific parts of the branches of the plant; (4) differential distribution in shaded and open areas; and (5) differential altitudinal distribution of the two *Peucetia* species. With respect to temporal niche, we observed that the two spider species had a similar age structure and similar fluctuation in abundance throughout the year. With respect to micro-habitat use, in both species, different instars used different plant parts, while the same instars of both species used the same type of substrate. However, the two *Peucetia* species segregated by meso-habitat type, with *P. rubrolineata* preferring *T. adenantha* plants in shaded areas and *P. flava* preferring those in open areas. Our results support the hypothesis of niche partitioning begetting diversity, and highlight the importance of analysing habitat use at multiple scales to understand mechanisms related to coexistence.

## INTRODUCTION

The mechanisms involved in species coexistence are a central theme in ecology, as they are responsible for maintaining high species diversity in ecosystems worldwide (Stevens 1989; Hubbell 2001; Tilman 2004; Siepielski & McPeek 2010: Brown 2014). Competitive interactions are stronger between morphologically similar and phylogenetically close sympatric species (Chillo et al. 2016). The first mathematical models of resource competition proposed that when two species compete for the same resource, one species invariably eliminated the other (Vance 1984). However, considerable species diversity can be observed coexisting and persisting in the same trophic level of a given web of species interactions (Schoener 1974; Kelly & Bowler 2009). Hutchinson (1957) suggested that the coexistence of ecologically similar species at the same site requires a form of resource partitioning.

Partitioning would enable coexistence by reducing or preventing interspecific competition. Resource use partitioning has been reported for several taxa, including, mammals (Chillo et al. 2010; Di Bitetti et al. 2010), birds (Young et al. 2010; Novcic 2016), reptiles and amphibians (Toft 1985; Segurado & Figueiredo 2007) fishes (Ross 1986), and invertebrates (Boyer & Rivault 2005). These studies showed that several mechanisms, such as differences in phenology or specific habitat selection, led to a decrease in competition, allowing the species to coexist.

One particular group whose coexistence mechanisms have been given considerable attention are spiders (Michalko & Pekár 2015). Spiders are one of the most diverse group of terrestrial predators (Turbull 1973; Wise 1993), with up to 4 species and 131 individuals co-occurring per m^2^ in tropical forests (Castanheira et al. 2016; Rodrigues et al. 2016; Nyffeler & Birkhofer 2017). Partitioning of resources among spiders can be related to: (1) variation in prey types or sizes as a consequence of different hunting strategies or body size (Olive, 1980; Nyffeler 1999; Yamanoi & Miyashita 2005; Novak et al. 2010); (2) activity time, with different species varying in phenologies or circadian cycles (Herberstein & Elgar 1994; Herberstein 1997; Novak et al. 2010); (3) use of space, with different species using different microhabitats as the result of cryptic colorations (Oxford & Gillespie 1998), different foraging behaviors or different sites chosen for the placement of webs (e.g., different heights) (Aiken & Coyle 2000, Ganier & Höfer 2001, Henaut et al. 2006, Harwood & Obrycki 2003, Cumming & Wesolowska 2004; Portela et al. 2013; Souza et al. 2015). Despite the variety of mechanisms mediating coexistence in spiders, only a few studies have assessed different niche dimensions simultaneously in sympatric and ecologically similar species (e.g. Gasnier & Höfer 2001, Michalko et al. 2016). Understanding how different partitioning mechanisms acting through multiple niche dimensions contribute to the coexistence of ecologically similar organisms can help us advance in the understanding of the processes shaping diversity patterns in local scales. Here we study different aspects of habitat use of two co-occurring spiders in the genus *Peucetia* (Oxyopidae). *Peucetia* is a cosmopolitan group, comprising 47 species, with most species occurring in tropical regions (Santos & Brescovit 2003; World Spider Catalog 2018). Spiders of this genus do not build webs; they weave wire guides between the branches, flowers, or leaves of plants where they live. Some *Peucetia* species were observed to be associated with more than 50 species of plants belonging to 17 families, all with glandular trichomes, showing their strong associations with this plant characteristic (Vasconcellos-Neto et al. 2007). The species *Peucetia rubrolineata* Keyserling, 1877 and *P. flava* Keyserling, 1877, both of similar sizes (13–14 mm) are often found in *Trichogoniopsis adenantha* shrubs (DC.) (Asteraceae) (King & Rob 1972) and appear to use the same micro-habitat and food resources (King & Rob 1972). They also have similar phenologies (Villanueva-Bonilla et al. 2018). Considering all ecological similarities, this pair of species living in the same plant comprises a promising study system to understand niche partitioning. In this sense the question we address here is: how do these species partition resources? We propose the following hypotheses to explain the coexistence of these two *Peucetia* species: (1) temporal segregation: the reproductive cycle of the species and the hatching of the eggs could occur in different months (different season on the year); 2) microhabitat segregation—stratification on plants: *P. rubrolineata* and *P. flava* occupy different types of branches (vegetative branches vs. reproductive branches) or different plant parts (e.g., leaves, stems, and flower heads). 3) mesohabitat segregation—patches with different environmental conditions: *Peucetia* species may occur preferentially in different environments with respect to luminosity and humidity. 4) macrohabitat segregation—altitudinal separation: *P. rubrolineata* and *P. flava* may be distributed differently along the altitude gradient in the Serra do Japi (740–1,294 m).

## MATERIALS AND METHODS

### Study area

This study was carried out at Serra do Japi, located between 23° 11’ S and 46° 52’ W of the Atlantic Plateau between the municipalities of Jundiaí, Itupeva, Cabreúva, Pirapora do Bom Jesus, and Cajamar in the state of São Paulo, Brazil,. Located between 700 m and 1,300 m elevation this area (354 km^2^) comprises seasonal mesophyllous forests lies. The climate is seasonally well defined, with average monthly temperatures varying from 13.5 °C in July to 20.3 °C in January and with a rainy season in summer (December–March) and dry season in winter (June–August) (Pinto 1992). Surveys were conducted in different regions of the Serra: (1) at the dam of the DAE (i.e., the dam of the department of water and sewage); (2) near the field base at the area locally known as “Biquinha”; (3) along a regional pathway at a locality named “TV cultura” at two altitudes (900–1000 m and 1170–1290 m). To determine the role of (1) time segregation, (2) microhabitat—stratification on plants, and (3) mesohabitat—patches with different environmental conditions in resource partitioning, studies were conducted from 2013–2015. To determine the role of macrohabitat—altitudinal distribution, research was carried out in March 2014 and March 2017.

### Time segregation

#### *Population dynamics of* Peucetia rubrolineata *and* Peucetia flava

Over two years, we recorded the fluctuation and phenology (age structure over time) of the *P. rubrolineata* and *P. flava* populations by monthly observations of these spiders on 100 and 200 plants, respectively, in shaded and open areas. For each individual, we recorded the species, instar, and the specific part of the plant where the spider was located. We identified the instars in accordance with the guidelines published by Romero and Vasconcellos-Neto (2005). This classification was based on the size of the cephalothorax and the first pair of legs (For details see Villanueva-Bonilla & Vasconcellos-Neto 2016; Villanueva-Bonilla et al. 2018).

### Spatial segregation

#### Microhabitat—stratification on plant

##### *Phenology of the plant* Trichogoniopsis adenantha

To verify whether the phenology of *T. adenantha* affects the occurrence, population dynamics, or age structure of *P. rubrolineata* and *P. flava*, each month, we recorded the number of vegetative branches and the developmental stage of the flower heads of 20 randomly selected plants.

For the phenological description of the flower heads, we classified them in situ into five phenophases as described by Almeida (1997) and Romero (2001): (F1) closed bud, very small, bracts covering the whole bud; (F2) open bud, with all flowers visible but all closed (pre-anthesis); (F3) open flowers with long and bluish-pink stigmas (anthesis and fertilization); (F4) complete formation of yellow flowers on flower head; stigmas beginning to drop (fruit development phase); and (F5) dry flower head; mature seeds, beginning of the dispersion phase.

Subsequently, based on the predominance of the phenophases of each flower head, we classified the branches into the following categories: (1) vegetative branches; (2) branches with flower buds, with the mean number of flowers in the F1 or F2 phase greater than 50% of the total number of flower heads (owing to inter-individual variation in the number of flower heads from 6 to 18 flower heads per branch); (3) branches with open flowers, with the mean number of flowers in the F3 greater than 50% of the total number of flower heads; (4) type 4 branches, with the mean number of flowers in the F4 greater than 50% of the total number of flower heads; (5) type 5 branches, with the mean number of flowers in the F5 greater than 50% of the total number of flower heads.

##### *Distribution of* Peucetia rubrolineata *and* Peucetia flava *instars on* Trichogoniopsis adenantha *branches*

To verify whether the distribution of the *P. rubrolineata* instars was similar in the different types of branches, we also recorded the frequency of the different types of branches available and those actually occupied by the spiders. For the analysis, only data from the months where *P. rubrolineata* was most abundant were used to ensure a sufficient numbers of individuals for analysis. We pooled the information of the instars of the four inspected localities for this analysis. The same procedure was performed for *P. flava*.

##### *Micro-site on the branch of the* Trichogoniopsis adenantha plant

We applied the Chi-square Test (see Sokal & Rohlf 1994; Zar 1998) to verify whether instars of *P. rubrolineata* have the same pattern of distribution in different specific parts of the plant (leaf, stem, and flower head and dry flower head). As the null hypothesis, we predicted the spiders of each instar would occur on different parts of the branch with similar probability. We also adopted the same protocol for *P. flava*.

##### *Similarity in the distribution of instars of the two* Peucetia *species on* Trichogoniopsis adenantha

Once the occurrence of each instar of each *Peucetia* species in the different parts of the plant (leaf, stem, flower head, dry flower head) was known, we investigated whether the same instars of both species occupy the same part of the plant using the Chi-squared Test to examine the significance of observed differences in the frequency of occupation. To do this, we related the total available parts of the inspected plants to the parts that were actually occupied by each instar of both *Peucetia* species.

#### Mesohabitat—patches with different environmental conditions

##### Plant habitat

We examined *T. adenantha* plants to identify both *Peucetia* species, totaling 100 individuals for each spider species. Subsequently, to determine the light volume gradient above each inspected plant, we placed a camera with a fisheye lens on each *T. adenantha* plant to photograph vegetative cover above it. We took photographs only on plants where *Peucetia* individuals were present. Subsequently, we divided photographs into quadrants and classified each site according to canopy cover percentage classes: 0– 20%; 21–40%; 41–60%; 61–80%, and 81–100%.

We compared the total frequency of each species in each vegetation cover class and used a Chi-squared Test to test whether the occupation patterns of the two spider species differed between the different classes of vegetation cover.

##### Co-occurrence of both spider species

During the two years of study, for each *T. adenantha* plant observed, we recorded the species of spider present and the number of individuals of each species. To determine the degree of niche overlap of the two species of *Peucetia* in each instar, an index ranging from 0 to 1 was defined, where 0 is no overlap and 1 is 100% overlap. To calculate the index of each instar, we put together the number of plants recorded in the two years of study in which the two species were present simultaneously, divided by the total number of plants that had at least one spider of any species of *Peucetia*. The index was calculated for the same instar of the two species of *Peucetia* separately because the two species presented in similar proportions of abundance throughout the year and the same instars of the two species occupy the same type of branch on the plant.

#### Macrohabitat—altitudinal separation

##### Altitudinal distribution

To verify whether *P. rubrolineata* and *P. flava* occur at different altitudes in the Serra do Japi, we conducted sampling in March 2014 in an altitudinal transect in three altitudinal bands: (1) 800–900 m, dam of the DAE; (2) 900–1,100 m, two sites, (a) base region, shaded area, and (b) regional pathway of “TV culture,” open region; (3) 1,170– 1,290 m regional “TV culture” pathway at different altitudes. In each altitudinal range, we inspected the *T. adenantha* plants in open and shaded areas: 199 plants (149 and 50 plants in shaded and open areas, respectively) in the range of 800–900 m; 189 plants (29 and 160 plants in shaded and open areas, respectively) at site (a) and 113 (98 and 15 plants in shaded and open areas, respectively) plants at site (b) at altitude between 900– 1,100 m; 115 plants (20 and 95 plants in shaded and open areas, respectively) at altitude of 1,170–1,290 m. The numbers of plants available for inspection varied at different altitudes. Regarding the altitude range of 900–1,100 m, in 2014, the location (a) was an open area because mixed vegetation of native trees and *Eucalyptus* had been cut and, consequently, *T. adenantha* plants were more abundant. Three years later (2017), the tree vegetation grew and shaded most of the *T. adenantha*. Thus, we sampled this area again in 2017 to verify the effect of shading on the relative abundance of the two *Peucetia* species. In each strip, we recorded the environment in which the plant was (open or shaded according to the photographs of canopy cover) and the *Peucetia* species found on each plant. We used an altimeter to measure altitude. Subsequently, we applied the Chi-squared test (with Williams’ correction) to compare the frequencies of the two spider species with the frequencies of *T. adenantha* in open or shaded environments at different altitudes. We conducted these analyses to separate the effect of altitude with respect to the occurrence of these spiders in different microenvironments, in the case of open areas and more shaded areas.

For all comparisons regarding the frequencies of use of habitat - temporal segregation, ontogenetic variation, microhabitat segregation and macrohabitat partitioning - we used a Chi-square test with Monte Carlo permutations with 10000 simulations (Hope 1986) to test for significance considering an alpha value of 0.05. In cases of multiple pairwise comparisons we used the Williamś correction. For all statistical analyzes, we used the free software R (R Core Team, 2016). The package “rmngb” was used for all Chi-squared analyses.

## RESULTS

### Temporal segregation

#### Population dynamics of Peucetia rubrolineata and Peucetia flava

A total of 726 individuals of *P. rubrolineata* and 628 of *P. flava* were observed during the two-years study. The population fluctuation and age structure of the two *Peucetia* species were similar throughout the study period. The two species showed similar temporal peaks of abundance (in summer), and similar patterns of appearance of different instars throughout the year (Fig. 1).

**Fig. 1.**
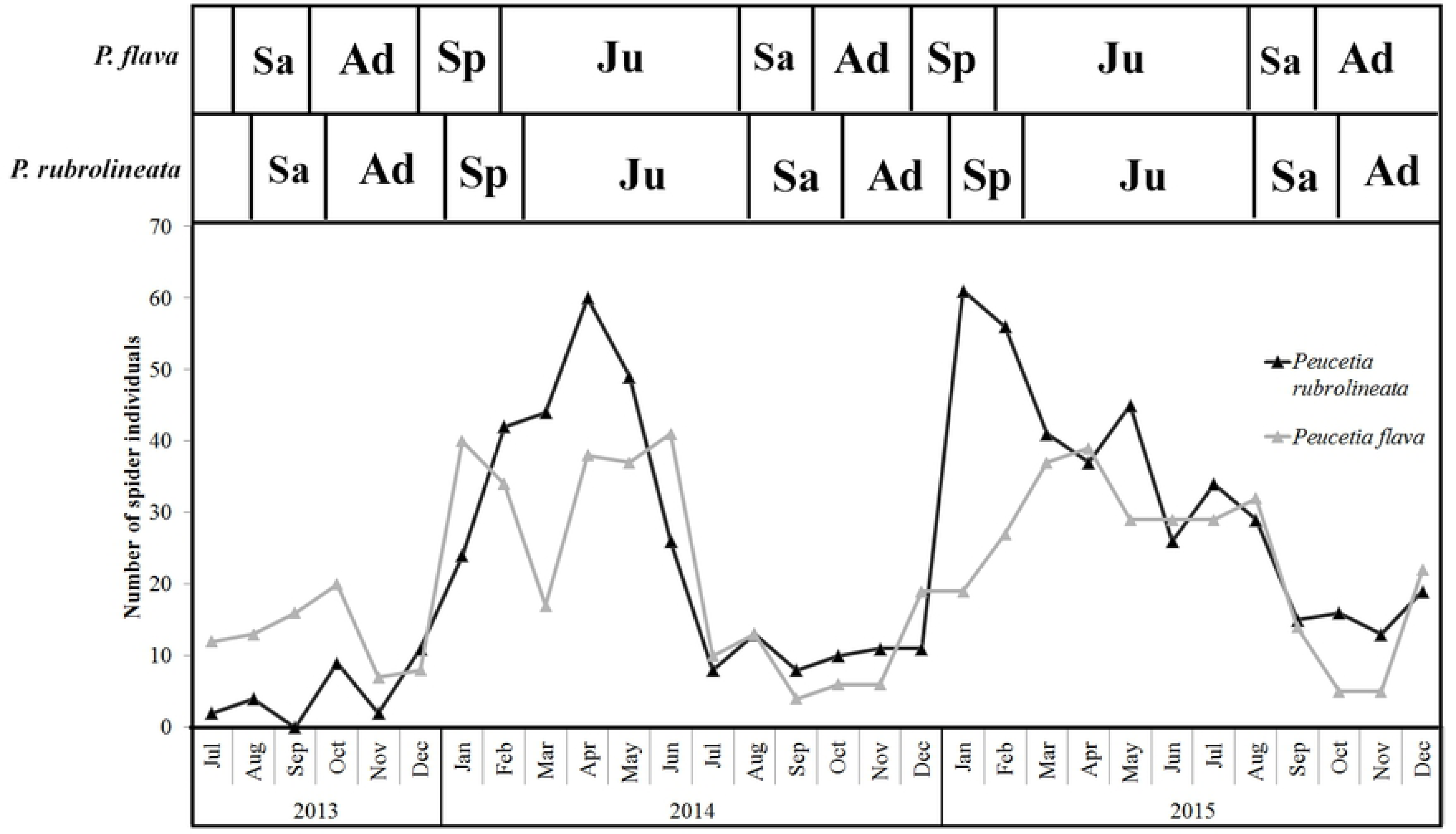
Population fluctuation of *Peucetia rubrolineata* and *P. flava* indicating periods of abundance (upper bars) of each of the developmental instars throughout the study period. **Sa** = subadult individuals; **As** = adult individuals; **Sp** = Spiderlingss; **Ju** = young individuals + juveniles individuals.

#### Phenology of Trichogoniopsis adenantha

In general, *T. adenantha* has both vegetative and reproductive branches throughout the year; however, the proportions vary with the season (Fig. 2). In winter and spring, vegetative branches occur in larger proportions. At the end of the dry season, usually in spring (months), there is growth of vegetative branches, followed by an increase in the number of reproductive branches, with a large number of buds in spring and summer. During March and April, a period with greater abundance of branches with open flowers occurs. In May, a large number of flowering branches occurs in phase 4 (seed formation), and in June and July, there are several flowering branches in phases 4 and 5 dry (flower heads). In general, vegetative branches were more abundant than reproductive branches. Production of flower heads occurred all over the year, however, the proportion of each phenophase varied from year to year (Fig. 2).

**Fig. 2.**
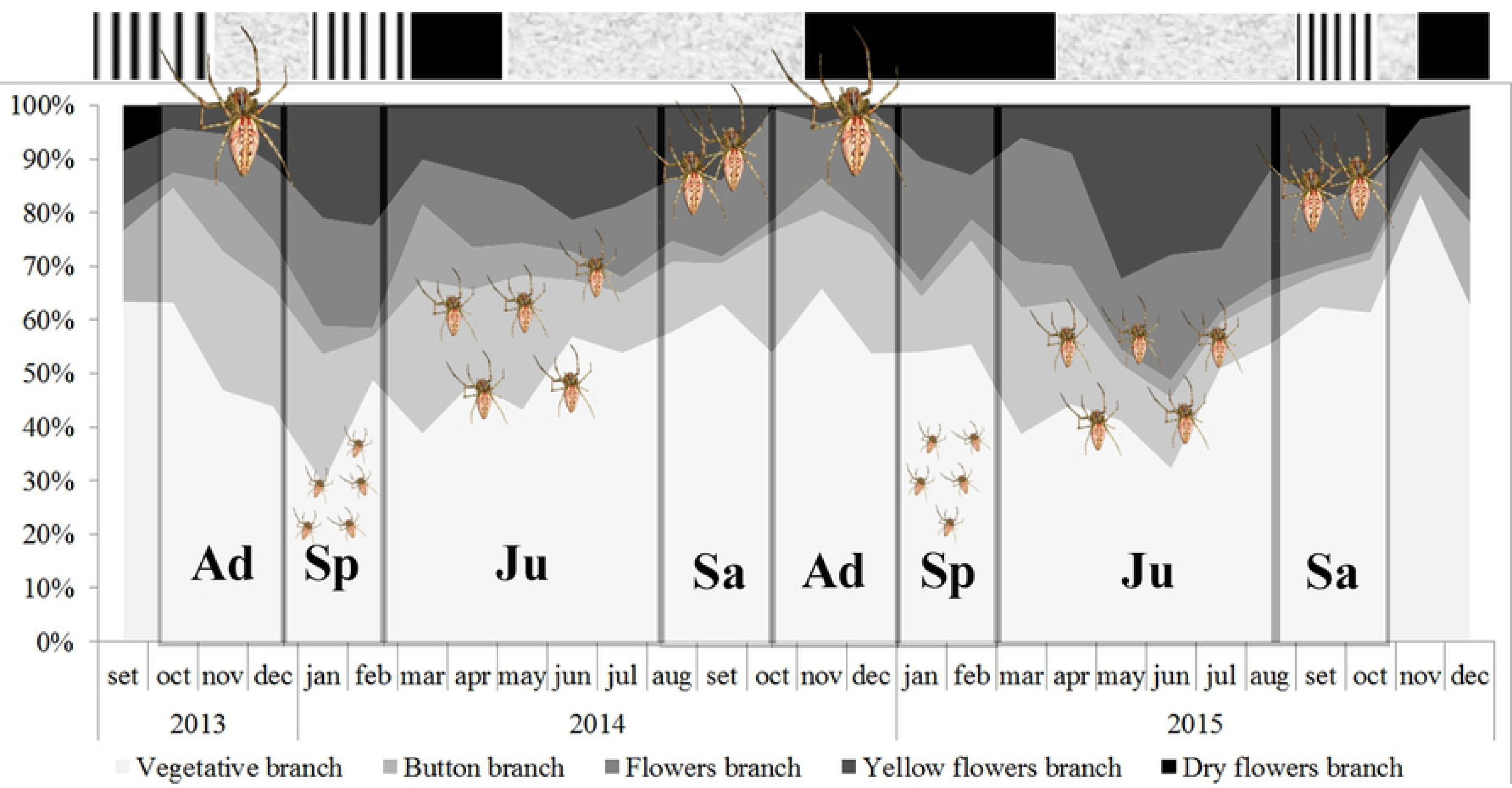
Phenogram of the different types of *T. adenantha* branches with the representation of the different age classes of *Peucetia* in the type of branch where they are frequently found. **Ad** = adult individuals; **Sp** = Spiderlings; **Ju** = Young and Juvenile; **Sa** = subadult individuals. The upper bar of the graph represents the rainy periods where: (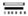) Dry period; (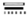) Period with average rainfall; (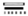) Rainy season.

### Spatial segregation

#### Microhabitat—stratification on the plant

##### Phenology of Trichogoniopsis adenantha and distribution of the Peucetia species on T. adenantha branches types

The distribution of each instar of the two *Peucetia* species across the types of branches did not follow the relative abundance of these branches throughout the year; the differential utilization of the branches by the two species was maintained the same regardless of branch proportion (Fig. 2). Thus, young spiders were more frequent in the vegetative branches (Chi-squared test= *P. rubrolineata*: X^2^ = 9.37, *P* = 0.034; *P. flava*: X^2^ = 10.777, *P* = 0.005; in 10000 simulations). Juvenile spiders were more frequent on vegetative branches and in 5 type branches, although the other types of branches are more frequent (Chi-squared test=*P. rubrolineata*: X^2^ = 8.31, *P* = 0.043; *P. flava*: X^2^ = 11.111, *P* = 0.0093). Subadult spiders were also more frequent on vegetative branches and in 5 type branches, whilst vegetative branches were more frequent (Chi-squared test= *P. rubrolineata*: X^2^ = 11.327, p = 0.0055; *P. flava*: X^2^ = 8.512, p = 0.0069). Adult individuals were more frequent on type 5 branches, especially in the inflorescences where shelters made with leaves and dry flower heads are often observed (Chi-squared test=*P. rubrolineata*: X^2^ = 10.096, *P* = 0.0038; *P. flava*: X^2^ = 12.791, *P* = 0.00089). On the other hand, spiderlings of the two *Peucetia* species were the only ones that presented similar observed and expected frequencies in all types of branches (Chi-squared test=*P. rubrolineata*: X^2^ = 4.22, *P* = 0.215; *P. flava*: X^2^ = 1.22, *P* = 1) (Fig 3).

**Fig 3.**
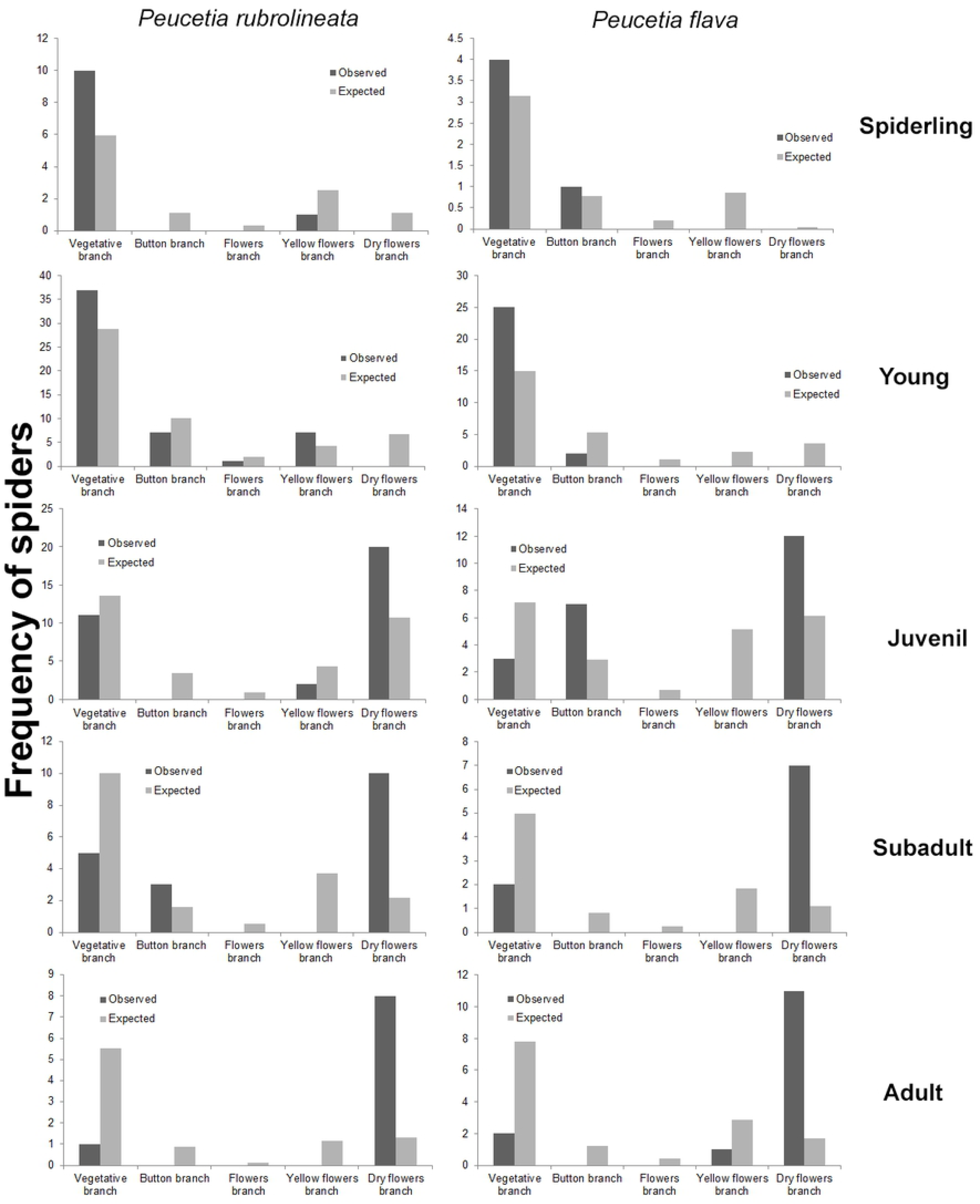
Observed and expected frequencies of the *P. rubrolineata* e *P. flava* instars on different types branches of *T. adenantha*.

##### Micro-sites on the branches of Trichogoniopsis adenantha plants

The distribution of different instars of *P. rubrolineata* and *P. flava* on different parts of the plant was not homogeneous (Table 1, 2).

**Table 1.**
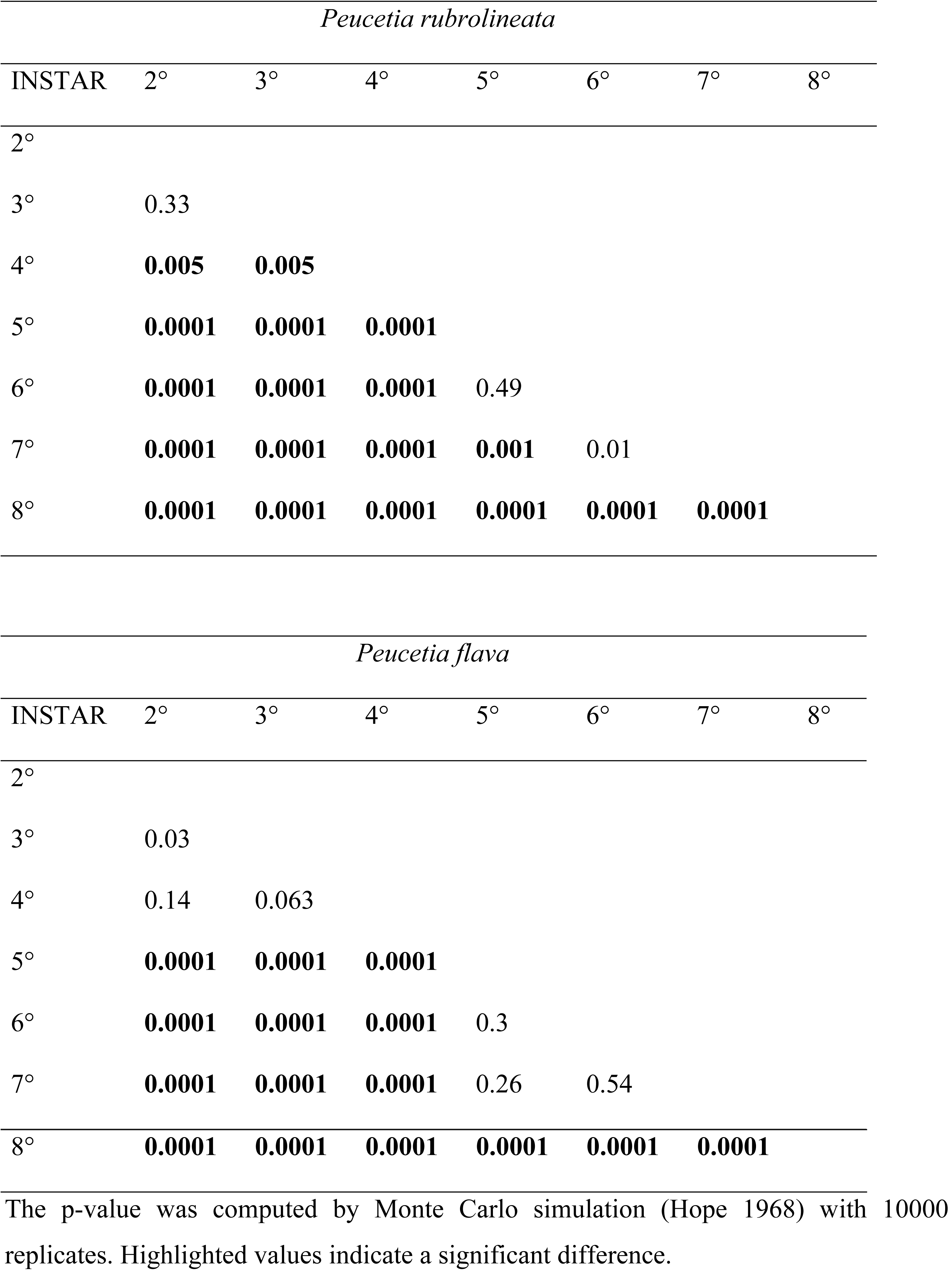
Comparison of the distribution of the developmental instars of each of the species of *Peucetia* on the different parts of the plant *Trichogoniopsis adenantha*.

| *Peucetia rubrolineata* |  |  |  |  |  |  |  |
| --- | --- | --- | --- | --- | --- | --- | --- |
| INSTAR | 2° | 3° | 4° | 5° | 6° | 7° | 8° |
| 2° |  |  |  |  |  |  |  |
| 3° | 0.33 |  |  |  |  |  |  |
| 4° | **0.005** | **0.005** |  |  |  |  |  |
| 5° | **0.0001** | **0.0001** | **0.0001** |  |  |  |  |
| 6° | **0.0001** | **0.0001** | **0.0001** | 0.49 |  |  |  |
| 7° | **0.0001** | **0.0001** | **0.0001** | **0.001** | 0.01 |  |  |
| 8° | **0.0001** | **0.0001** | **0.0001** | **0.0001** | **0.0001** | **0.0001** |  |
| *Peucetia flava* |  |  |  |  |  |  |  |
| INSTAR | 2° | 3° | 4° | 5° | 6° | 7° | 8° |
| 2° |  |  |  |  |  |  |  |
| 3° | 0.03 |  |  |  |  |  |  |
| 4° | 0.14 | 0.063 |  |  |  |  |  |
| 5° | **0.0001** | **0.0001** | **0.0001** |  |  |  |  |
| 6° | **0.0001** | **0.0001** | **0.0001** | 0.3 |  |  |  |
| 7° | **0.0001** | **0.0001** | **0.0001** | 0.26 | 0.54 |  |  |

|  |  |  |  |  |  |  |
| --- | --- | --- | --- | --- | --- | --- |
| 8° | 0.0001 | 0.0001 | 0.0001 | 0.0001 | 0.0001 | 0.0001 |

The p-value was computed by Monte Carlo simulation (Hope 1968) with 10000 replicates. Highlighted values indicate a significant difference.

**Table 2.** Abundance of spiders of *Peucetia rubrolineata* and *P. flava* (Oxyopidae) in different parts of the plant *Trichogoniopsis adenantha* (Asteraceae).

| *Peucetia rubrolineata* |  |  |  |  |  |  |  |  |
| --- | --- | --- | --- | --- | --- | --- | --- | --- |
| Instar | Leaf | Steam | flower heads | Dry flower heads | Shelter | Total | X<sup>2</sup>-test | P value |
| 2° | 21 | 5 | 1 | 2 | 0 | 29 | 19.72 | **0.0003** |
| 3° | 84 | 31 | 22 | 14 | 0 | 151 | 62.78 | **0.00009** |
| 4° | 49 | 22 | 25 | 30 | 0 | 126 | 33.46 | **0.0009** |
| 5° | 47 | 9 | 23 | 111 | 0 | 190 | 96.3 | **0.0009** |
| 6° | 25 | 5 | 12 | 79 | 1 | 122 | 67.42 | **0.0009** |
| 7° (subadult) | 10 | 13 | 4 | 34 | 1 | 62 | 24.32 | **0.0002** |
| 8°(adult) | 8 | 1 | 0 | 10 | 37 | 56 | 34.13 | **0.0009** |
| *Peucetia flava* |  |  |  |  |  |  |  |  |
| Instar | Leaf | Steam | flower heads | Dry flower heads | Shelter | Total | X<sup>2</sup>-test | P value |
| 2° | 10 | 8 | 6 | 0 | 0 | 24 | 12.36 | **0.0061** |
| 3° | 49 | 21 | 6 | 9 | 0 | 85 | 40.65 | **0.0009** |
| 4° | 42 | 28 | 20 | 15 | 0 | 105 | 30.02 | **0.0009** |
| 5° | 32 | 8 | 18 | 91 | 0 | 149 | 76.36 | **0.0009** |
| 6° | 18 | 6 | 14 | 77 | 2 | 117 | 60.01 | **0.0009** |
| 7° (subadult) | 11 | 1 | 3 | 31 | 1 | 47 | 28.54 | **0.0009** |
| 8°(adult) | 15 | 1 | 3 | 20 | 50 | 89 | 41.20 | **0.0009** |

Highlighted values indicate significant values.

Second and third instars of *P. rubrolineata,* were more often recorded on the leaves, whereas fourth-instar spiders were observed on the leaves, stem, and flower heads, the fifth, sixth, and seventh instars occurred more frequently in the dry flower heads, and the eighth-instar spiders (adults) occurred more frequently in dry flower heads and in shelters made from remnants of the dry flower heads. *P. flava* individuals of the second, third, and fourth instars were found in higher frequencies on the leaves of *T. adenantha*; fifth, sixth, and seventh instars also occurred more frequently in the dry flower heads, and adult spiders (eighth instar) also occurred more frequently in dry flower heads and shelters (Fig. 4–5).

**Fig. 4.**
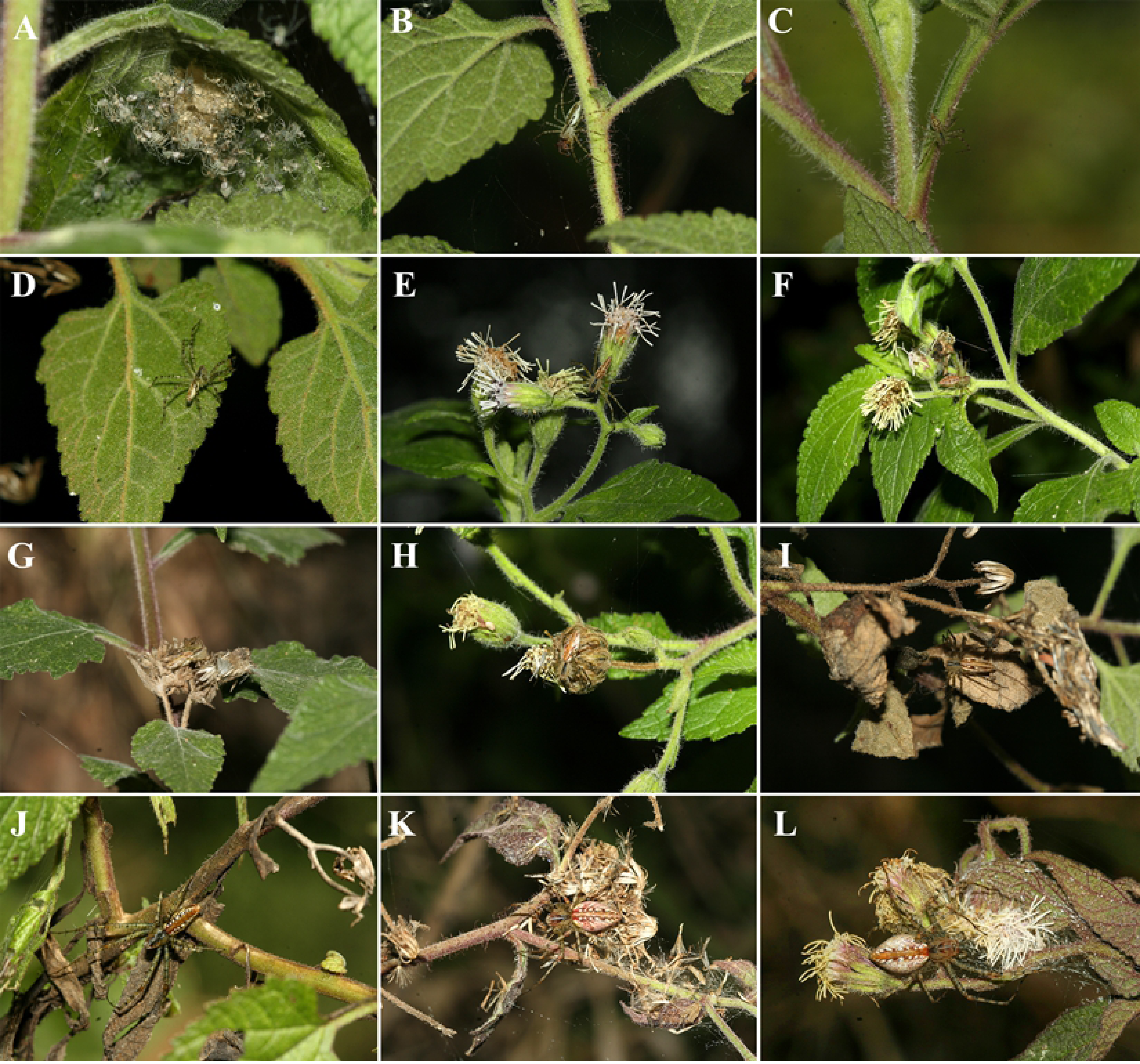
Instars of *Peucetia rubrolineata* and *P. flava* on different parts of the plant *Trichogoniopsis adenantha* (Asteraceae) where they are frequently observed (leaf, stem, flower heads, flower, shelter of dry flower heads). A = second instar *P. rubrolineata*; B-D = third instar *P. rubrolineata*; E = fourth instar *P. rubrolineata*; F = fourth instar *P. flava*; G-H = fifth instar *P. rubrolineata*; I = sixth instar *P. rubrolineata*; J = seventh instar (subadult) *P. flava*; K-L = eighth instar (adult) *P. flava*.

**Fig. 5.**
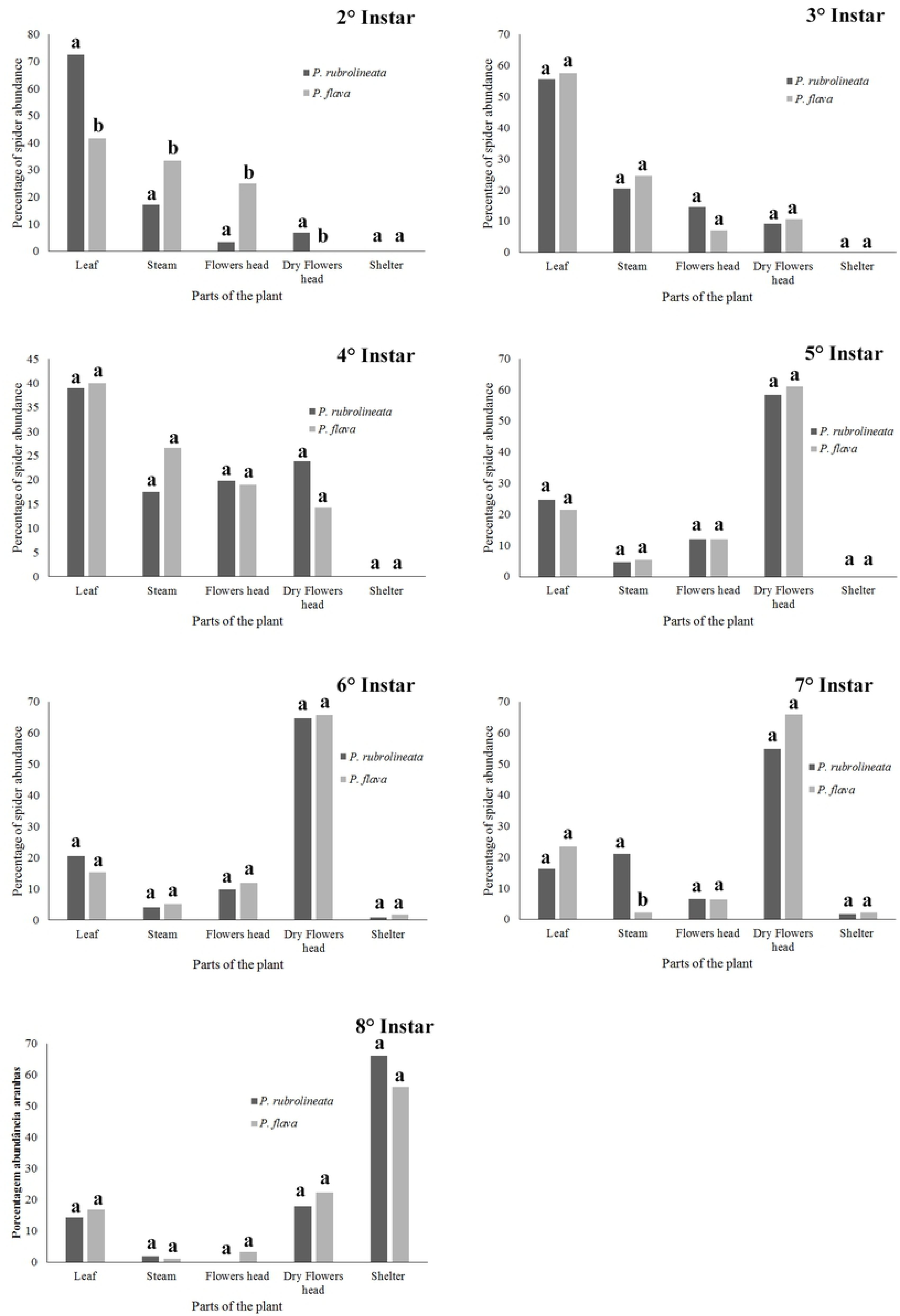
Comparison of the observed frequency of developmental instars of Peucetia *rubrolineata* and *P. flava* (Oxyopidae) on the different parts of the plant *Trichogoniopsis adenantha* (Asteraceae) in Serra do Japi, SP. Brazil. Instar 7° = subadult; Instar 8 ° = adult. Different letters indicate statistically different values.

Despite the difference in the occupation of the types of branches and parts of the plant between instars of the same species when the same instars of the two species are compared, the pattern of occupation on the plant is similar. (Fig. 5, Table 2).

#### Mesohabitat—patches with different environmental conditions

##### Co-occurrence of both Peucetia species

The frequencies of the two species of *Peucetia* were different in the two types of vegetation cover (X^2^ = 75.444, *P* = 0.00009; in 10000 simulations) (Fig. 6). *P. rubrolineata* was recorded at a higher frequency in *T. adenantha* plants located in more close canopy environments (X^2^ = 41.622, *P* = 0.00009) (Fig. 6B). In contrast, *P. flava* was recorded more frequently in *T. adenantha* plants in more open environments (X^2^ =13.085; *P* = 0.0102) (Fig. 6A). In the other vegetation cover types, the frequencies of both *Peucetia* species were similar (Fig. 6C).

**Fig. 6.**
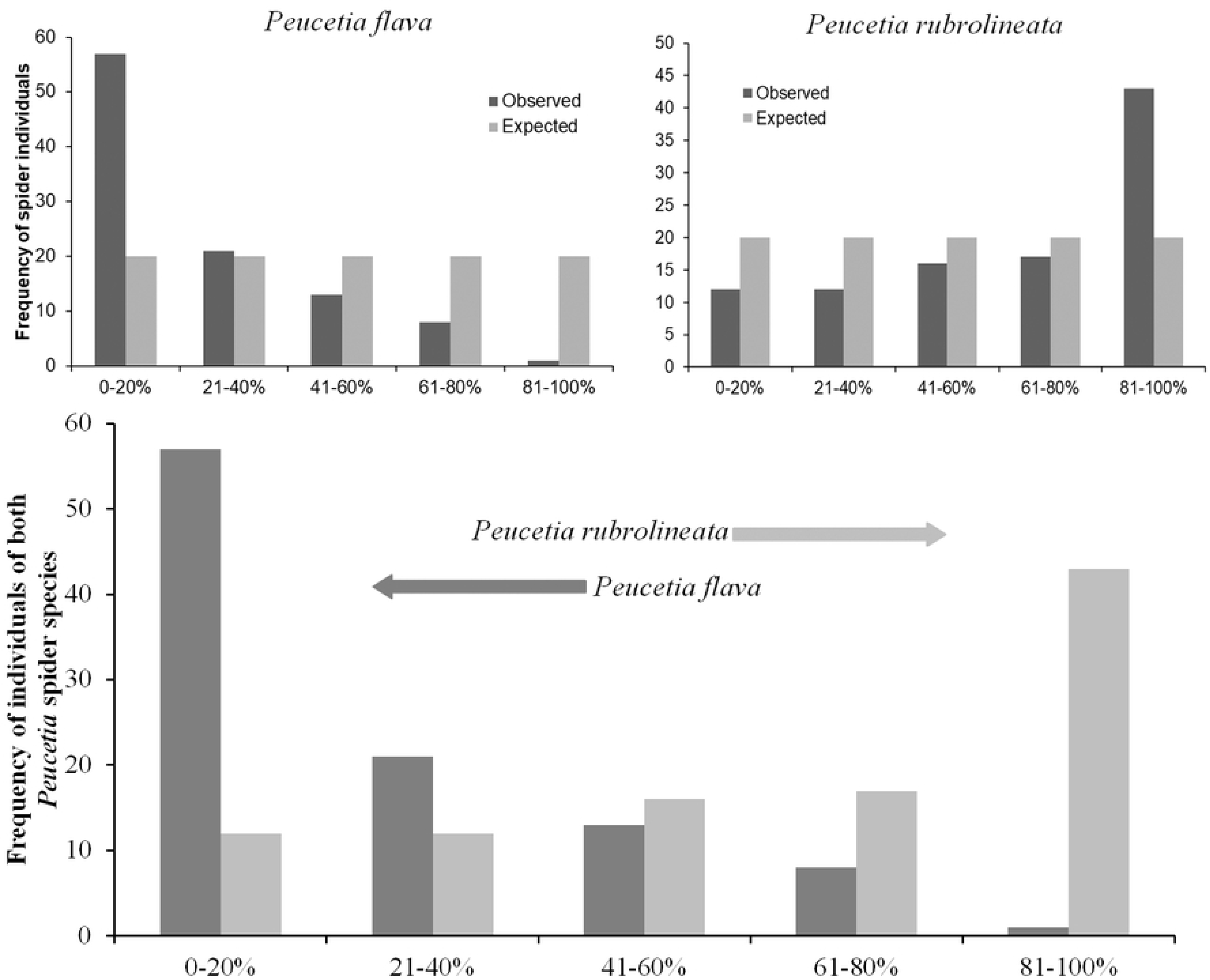
Distribution of *Peucetia rubrolineata* and *P. flava* on *Trichogoniopsis adenantha* plants in environments with different vegetation cover (%). Observed and expected frequencies of A) *P. flava* and B) *P. rubrolineata* in different percentages of vegetation cover.

In environments with intermediate canopy cover, a larger number of plants were observed to harbor the two *Peucetia* species. Of the 729 plants that had spiders during the study period, 43.5% contained only *P. flava* and 53.3% contained only *P. rubrolineata*, whereas only 3.4% of the plants contained both species, indicating little overlap of their niches.

The values of niche overlap varied from 0.024 to 0.059, with a mean of 0.034, indicating low overlap between the same instar of the two species of *Peucetia* (Table 3).

**Table 3.** Niche overlap index between the same instar of the two species of *Peucetia* and the average value of niche overlap.

| Instars | Total plants<br>with spiders | Plants with two<br>spider species | Niche overlap<br>index |
| --- | --- | --- | --- |
| 2<sup>nd</sup> | 36 | 0 | 0 |
| 3<sup>rd</sup> | 163 | 4 | 0.024 |
| 4<sup>th</sup> | 108 | 4 | 0.037 |
| 5<sup>th</sup> | 135 | 8 | 0.059 |
| 6<sup>th</sup> | 113 | 3 | 0.026 |
| 7<sup>th</sup> | 77 | 2 | 0.025 |
| 8<sup>th</sup> | 97 | 4 | 0.041 |
| Average value |  |  | 0.034 |

#### Macrohabitat—altitudinal separation

##### Altitudinal distribution

Both *Peucetia* species were found at all altitudes at the study site, but in different frequencies (Total *P. rubrolineata* 63 individuals; Total *P. flava* 37 individuals). However, these frequencies varied depending on availability of *T. adenantha* plants in more open areas (little canopy cover) or in more closed places (abundant canopy cover) and not with respect to the altitude. Sites with low canopy cover harbored individuals of *P. flava,* whereas sites with abundant canopy cover harbored individuals of *P. rubrolineata* (Table 4). In the observation made in 2014 at site (2b), called the TV Cultura Pathway Region, *T. adenantha* plants were found with little canopy cover and a higher abundance of *P. flava* was recorded. Three years later, in 2017, a higher abundance of *P. rubrolineata* was observed (Table 4) in the same region and at the same altitude (2b*), when vegetation grew and canopy cover increased on *T. adenantha* plants.

**Table 4.** Comparison of the frequencies of *P. rubrolineata* and *P. flava* on *Trichogoniopsis adenantha* in shaded and open areas.

| Altitude m | Plants in Shaded areas | Plants in open areas | Total of plants | Plants with *P. rubrolineata* | Plants with *P. flava* | X<sup>2</sup> Test | P value |
| --- | --- | --- | --- | --- | --- | --- | --- |
| (1) 800-900 | 149 | 50 | 199 | 20 | 5 | 0.179 | 0.74 |
| (2a) 900 -1100 | 98 | 15 | 113 | 13 | 4 | 0.594 | 0.64 |
| (2b) 900 -1100 | 29 | 160 | 189 | 1 | 13 | 0.473 | 0.59 |
| (2b*) 900 -1100 | 95 | 30 | 125 | 27 | 9 | 0.009 | 1 |
| (3) 1170 -1290 | 20 | 95 | 115 | 2 | 6 | 0.13 | 1 |

(1) DAE dam, (2a) Region of Base, more shaded, record made in 2014. (2b) Region path of TV culture, more open, record made in 2014. (2b\*) Same Region path of TV culture, record made in 2017 (more shaded), (3) Region path of TV culture at highest altitude, record 2014.

## DISCUSSION

Niche theory suggests that the coexistence of ecologically similar species requires a form of resource partitioning that reduces or prevents interspecific competition (Chesson 2000). In the studied system, the two *Peucetia* species inhabiting plants of *T. adenantha* presented similar phenology and population dynamics. In addition, the same instars of the two spider species had similar distributions across the different parts of the plant (i.e., similar use of the micro-habitat), indicating that the microhabitat overlap. However, our hypothesis that the two *Peucetia* species inhabiting *T. adenantha* plants are segregated has been confirmed, because they were distributed differently in response to shading (mesohabitat level). *Peucetia rubrolineata* occurred more frequently in places where canopy cover was higher, whereas *P. flava* occurred more frequently in more open sites. This pattern of distribution is probably affected by distinct physiological tolerance; however, further research is needed to verify this assumption. Results also indicated that these species have different altitudinal distributions because *P. rubrolineata* was more abundant at lower altitudes in the Serra do Japi, whereas *P. flava* occurred more frequently at higher altitudes. However, this difference seems to be more related to the availability of plants in different environments (shaded and sunny) than the altitude. At higher altitudes, the frequency of plants in the sunny areas was higher, as was the frequency of *P. flava*. In contrast, at lower altitudes, shaded areas were more frequent and the frequency of *P. rubrolineata* was higher. At intermediate altitudes, the abundance of spiders depended on the abundance of plants in the sun and shade. At the same site, the abundance of *P. flava* and *P. rubrolineata* varied in different years depending on the degree of shading. Although the two *Peucetia* species occurred in the intermediate shading environment, the co-occurrence of these spiders in the same host plant was infrequent.

*Peucetia rubrolineata* and *P. flava* showed no temporal segregation; both species exhibited population fluctuation and similar age structure throughout the year. In the literature, there is controversy over temporal segregation being one of the key mechanisms of coexistence in spider communities. On the one hand, Turner and Polis (1979) and Uetz (1977) suggested that temporal segregation is an important factor that reduces niche overlap. Gasnier and Höfer (2001) also argued that temporal segregation may facilitate the coexistence of species because the peak of abundance of each population will occur at different times, which reduces interspecific competition. For example, the spiders *Meta menardi* and *Metellina merianae* (Tetragnathidae), in addition to presenting distinct abundances throughout the year, also present distinct hunting strategies; *M. merianae* combines foraging outside and inside the web whereas *M. menardi* feeds exclusively on prey that fall into the web (Novak et al. 2010). Temporal segregation may also occur when species differ in periods of activity during the day (Krumpálová & Tuf 2013). Here we found no evidence that the studied species have temporal segregation related to population dynamics or individual behavior

Although *P. rubrolineata* and *P. flava* are sympatric and their population dynamics are similar throughout the year, the general pattern of spatial segregation registered occurred in the two years of study. This type of segregation as a central axis in the coexistence of sympatric spiders has been recorded in other cursorial spiders of the genus *Syspira* (Miturgidae), where two species of Miturgidae also exhibited different spatial distribution (differences in temperature and humidity at the site (Nieto-Castañeda & Jiménez-Jiménez 2009). In a central Amazon forest, Gasnier & Höfer (2001) also recorded significant differences in the habitat types used by four species of *Ctenus* (Ctenidae). In family Lycosidae, horizontal stratification appears to be a common mechanism of niche differentiation (Suwa 1986). In the temperate region the relative abundance of *Pardosa alacris* gradually increased with the canopy opening, whereas the opposite trend was observed for *P. lugubris*. Thus, niche differentiation along the canopy opening gradient mediated the coexistence of these two species in meta-communities (Michalko et al. 2016). The Lycosids *Geolycosa xera archboldi* and *G. hubbelli* also exhibited preferences for different habitats and micro-habitats based on the percentage of litter cover (Carrel 2003). All these examples show that habitat segregation at the mesohabitat level, as we found here, may be an important mechanism maintaining the high diversity of spiders in forested habitats.

Spiderlings of the two *Peucetia* species were more frequent on the vegetative branches of *T. adenantha*. A number of predatory arthropods (e.g., spiders) are specifically associated with glandular trichome-bearing plants (e.g., *T. adenantha*) where they capture prey attached to these sticky structures (Romero & Vasconcellos-Neto 2003; Vasconcellos-Neto et al. 2007). Spiderlings are probably more frequent in plant parts with more trichomes, which would increase the number of prey trapped (Romero et al. 2008). Moreover, trichomes provide some protection to the spiderlings from their enemies, such as other spiders (Langelloto & Denno 2004). Fourth-instar young spiders were observed preying on endophytic herbivores of the flower head of *T. adenantha*, especially *Trupanea* sp. (Tephritidae), and juveniles (fifth and sixth instars) predating larvae of Geometridae that feed on flower heads and other insects that inhabit the plant (J. Vasconcellos-Neto pers. Obs and see Romero & Vasconcellos-Neto 2005). In the case of subadults and adult spiders, they occurred more frequently in the flower head of the plant, especially those branches that also have dry structures where they rest. This higher frequency likely occurred because, besides having a place for camouflage, it is also easier for them to capture larger prey, such as floral visitors, which visit flower heads with open flowers at phase 3 and phase 4. In fact, Romero et al. (2008) recorded adult spiders feeding on *Pseudoscada erruca*, *Aeria olena*, and *Episcada carcinia* (Ithomiinae), and all floral visitors on the flowers of *T. adenantha*. Preferences for types of branches could be partially explained by the types of prey available and used as food by different instars. This ontogenetic habitat segregation found in both of the studied species reduces intraspecific overlap in resource use and may be important for the success of individuals at the different stages, and the population as whole.

By examining niche partition among two sympatric, morphologically similar, closely-related spiders at multiple hierarchically-organized spatial we show that despite the ecological similarity among these species, there is inter- and intraspecifc segregation in habitat use at different spatial scales. Although *P. flava* and *P. rubrolineata* are sympatric occurring in the same area with similar phenologies, at similar altitude, using the same parts of the same plants species, the distribution of the *T. adenantha* plant in shaded and open environments affects the distribution of the two species. Our results support the hypothesis of niche partitioning begetting diversity, and highlight the importance of analysing habitat use at multiple scales to understand mechanisms related to coexistence.

## ACKNOWLEDGEMENTS

This study was financed in part by the Coordenação de Aperfeiçoamento de Pessoal de Nível Superior - Brasil (CAPES) - Finance Code 001 (G.A. Villanueva-Bonilla). We thank the Prefeitura Municipal de Jundiaí, SP, Brazil for granting permission to work at the Biological Reserve of the Serra do Japi. The authors thank Instituto de Estudos dos Himenóptera Parasitóides da Região Sudeste Brasileira (INCT HYMPAR – SUDESTE-CNPq, FAPESP, CAPES).

## DISCLOSURE STATEMENT

No potential conflict of interest was reported by the authors.

